# Identification of TMEM230 mutations in familial Parkinson’s disease (response to comments)

**DOI:** 10.1101/170852

**Authors:** Han-Xiang Deng, Teepu Siddique

**Affiliations:** Ken and Ruth Davee Department of Neurology, Northwestern University Feinberg School of Medicine, Chicago, IL 60611, USA

## Abstract

We recently reported mutations in TMEM230 in familial Parkinson’s disease (PD). Farrer et al raised the concern that mutations in TMEM230 may not be pathogenic to PD. We seriously evaluated Dr. Farrer’s assertions. We obtained updated clinical information and performed several new experiments, including MegaEx chip screening of the family DNA samples with ∼2 million SNPs for whole-genome linkage study and re-analysis of whole-exome sequencing data. We did not find any other locus more robust than the chromosome 20p (TMEM230), nor any other variants with better segregation than TMEM230-R141L to explain the inheritance of PD in the large Mennonite family. Based on the new genetic data from the Mennonite PD family, and the robust genetic data showing additional TMEM230 mutations in multiple PD families, we are confident to conclude that *TMEM230* is a new PD-causing gene. Further studies of *TMEM230* should provide important mechanistic insights into understanding the vesicle/endosome trafficking/recycling defects in the pathogenesis of PD.

---

We recently reported the *TMEM230* c.422G>T (p.Arg141Leu, R141L) mutation in a large Mennonite PD family in Saskatchewan, Canada, a family that we had studied since 1992. Subsequently, we identified three additional mutations in the same gene, including a c.275A>G (p.Tyr92Cys, Y92C) in a sporadic PD case, a c.551A>G (p.*184Trpext*5) in a familial PD case, and a c.550_552delinsCCCGGG (p.*184ProGlyext*5) mutation in nine cases from seven PD families^1^. The Y92C and *184Trpext*5 were found in North America PD cases, and the *184ProGlyext*5 was identified in Chinese PD cases. Notably, both *184Trpext*5 and *184ProGlyext*5 occur at the stop codon, with an addition of five identical amino acids (HPPHS) at the C terminus of TMEM230 protein. These four mutations were neither present in our directly sequenced controls, nor in any public databases, including ExAC. Thus, our genetic findings are not restricted to a single Mennonite family, nor to a single ethnic population ^1^.

The same Mennonite PD family was also studied by Dr. Farrer’s group since 2010, but they reported a different mutation, *DNAJC13* c.2564A>G (p.Asn855Ser, N855S) in this family in 2014^2^. However, the DNAJC13-N855S mutation was not present in three affected cases, including two with PD (III-1 and III-23), and one with progressive supranuclear palsy (PSP) (II-1). Notably, the PSP case (II-1) was the father of a PD patient (III-1) without the N855S mutation. They argued that PSP might not be etiologically related with PD, and the two PD patients without N855S might be PD phenocopies ^2^. On inspection of the pedigree, because the father, five siblings and a son of the PSP case developed PD, the obvious explanation is that the PSP case is either an obligate carrier for the mutant PD gene or his PSP is a pleomorphism of the effect of the responsible gene mutation.

Subsequent to the publication of our paper, Dr. Farrer argued that genetic evidence for *DNAJC13* in PD remains greater than that for *TMEM230*, and made the assertions that the TMEM230-R141L mutation is not the genetic cause of PD in the large Mennonite PD family, and that TMEM230 is not a PD causing gene (http://biorxiv.org/content/early/2017/01/01/097030).

We seriously evaluated Dr. Farrer’s assertions, but did not find them substantiated by the data. To evaluate Dr Farrer’s concerns, we obtained updated clinical information and performed several new experiments, including MegaEx chip screening of the family DNA samples with ∼2 million SNPs. Dr. Farrer felt that the affection status of two individuals in the pedigree submitted by us was not accurate. Our affection status assignment was largely based on information from 2009 or prior. To update the affection status in this family, we contacted Dr. Ali Rajput, a PD specialist, who had originally ascertained this PD family. He reviewed his records and updated the affection status of the family members. Individual III-14, but not individual III-15, is affected. However, the individual III-14 has very mild and very slowly progressive symptoms starting age 59, but a diagnosis of PD could not be made in over a decade after the initial signs. He is now 82 years old. After 23 years of disease progression, he is still not on L-DOPA. This is the only affected case without TMEM230-R141L mutation, and likely represents a phenocopy. Whereas, there are three affected cases without DNAJC13-N855S mutation (II-1, III-1 and III-23). In addition, new linkage data from MegaEx chip screening containing ∼2 million SNPs consistently support the chromosome 20p (TMEM230) locus, which is consistent with our previous linkage data using genome-wide microsatellite markers; We did not find consistent whole-genome linkage data to support the *DNAJC13* locus or any other loci through our new studies using ∼2 million SNPs. Further analysis of exome sequencing and Sanger sequencing data showed that except for TMEM230-R141L, the other variants (including DNAJC13-N855S) were absent in at least three affected individuals in this family, and therefore, TMEM230-R141L remains the best genetic explanation for PD in this family (Table 1).

**Table 1.** Co-segregation analysis of variants shared by II-4 and III-26

| Individual<br>(Deng<br>et al) | Individual<br>(Vilarino-Guell<br>et al) | Pathology | AAO | TMEM230-R141L | DNAJC13-N855S | ZBTB38-R264Q | KIAA1109-H2613R | ADAMTS<br>L1-T128K | LGALS8-W359R | ABCB10-V321M | MS4A12-T202I | P2RX3-I26L | AGAP3-T284A | FAM13B-G496E | IQCD7-A7T | WDR17-V857M |
| --- | --- | --- | --- | --- | --- | --- | --- | --- | --- | --- | --- | --- | --- | --- | --- | --- |
| II-4 | II-1 | LB | 85 | + | + | + | + | + | + | + | + | + | + | + | + | + |
| II-6 | II-7 | LB | 68 | + | + | + | - | - | - | - | - | - | + | - | + | - |
| II-11 | II-11 |  | 80 | + | + | + | + | - | + | + | - | - | - | + | - | + |
| II-14 | II-9 | LB | 76 | + | + | + | - | + | + | + | + | + | + | + | + | + |
| III-16 | III-5 |  | 48 | + | + | + | - | + | + | + | + | + | - | - | - | - |
| III-17 | III-7 |  | 65 | + | + | + | - | + | - | - | + | + | - | - | - | - |
| III-18 | III-14 |  | 59 | + | + | + | - | - | + | + | - | - | - | - | - | + |
| III-20 | III-15 |  | 70 | + | + | + | - | - | - | - | - | - | - | + | - | + |
| III-26 | III-12 |  | 54 | + | + | + | + | + | + | + | + | + | + | + | + | + |
| III-23 | III-10 |  | 68 | + | - | - | + | - | + | + | - | - | - | - | + | + |
| III-14 | III-4 |  | 59 | - | + | + | + | + | + | + | - | + | - | - | - | - |
| II-1 | II-3 | PSP | 61 | + | - | - | + | + | - | - | - | - | - | + | + | - |
| III-1 | III-2 |  | 49 | + | - | - | + | + | - | - | - | - | - | - | - | - |

Note: Available DNA samples from 13 affected members were Sanger-sequenced for co-segregation of 13 variants identified by exome sequencing. These 13 variants are shared by II-4 and III-26. Because Vilarino-Guell et al used III-14 (the only phenocopy case without TMEM230-R141L mutation among all affected members in this family) for whole exome sequencing, the TMEM230-R141L was missed as a candidate pathogenic mutation. LB, Lewy body; PSP, progressive supranuclear palsy; AAO, age at onset.

The key data supporting the pathogenicity of *TMEM230* mutations, rather than the other variants, are summarized below.

1. Our new genome-wide linkage data using MegaEx chip screening containing ∼2 million SNPs demonstrated the most consistent high LOD scores in all models at the chromosome 20p (*TMEM230*) locus. The 20p (*TMEM230*) locus yielded the highest two-point and multi-point LOD scores, as shown in the affected-only model in the whole-genome. These new data are consistent with our previous linkage data using the whole-genome microsatellite markers ^1^. We did not find any other locus or loci better than 20p through our new whole-genome linkage analysis (Figs. 1-4, Table 2).
2. We updated the clinical information and re-analyzed our exome sequencing data (Table 1). TMEM230-R141L remains to be the most parsimonious explanation of etiology for PD in this family, as this mutation can explain the disease status of the entire family with a single phenocopy (II-14). DNAJC13-N855S and ZBTB38-R264Q require three phenocopies (II-1, III-1 and III-23), and the other mutations require five or more phenocopies (Table 1).
3. Although identification of the TMEM230-R141L mutation in the large Mennonite PD family was the original driving force leading us to test *TMEM230* mutations in other PD patients, we believe that our subsequent identification of two different mutations (*184Trpext*5 and *184ProGlyext*5) with nearly identical functional impact in 10 familial PD patients from 8 families in two well-separated populations (North America and China) **alone** is **sufficient** genetic evidence for the pathogenicity of *TMEM230* mutations to PD, regardless of the disease mechanism.

**Fig. 1.**
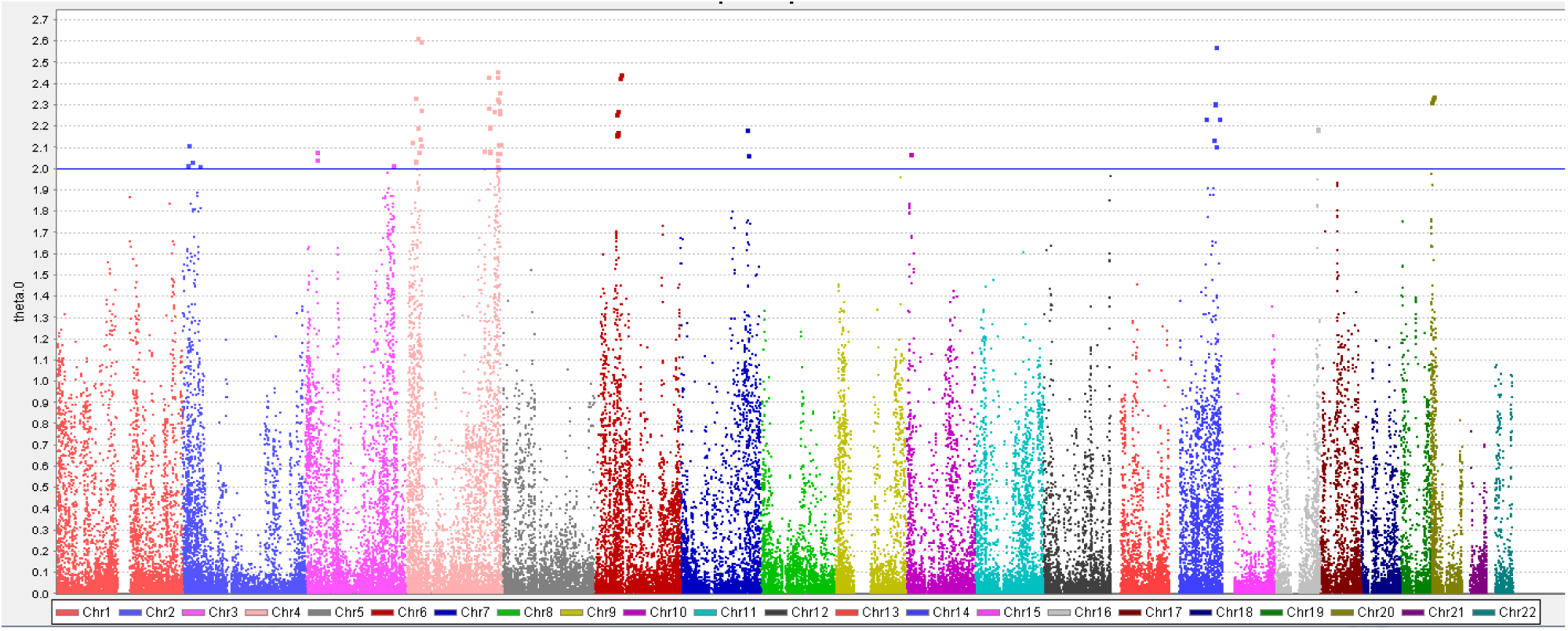
Genome-wide Fastlink two-point LOD scores for age-dependent penetrance model at theta=0 using MegaEx chip screening containing ∼2 million SNPs. Chromosomes 1-22 are color-coded. The positions of chromosomes are arranged from pter on the left side to qter on the right side.

**Fig. 2.**
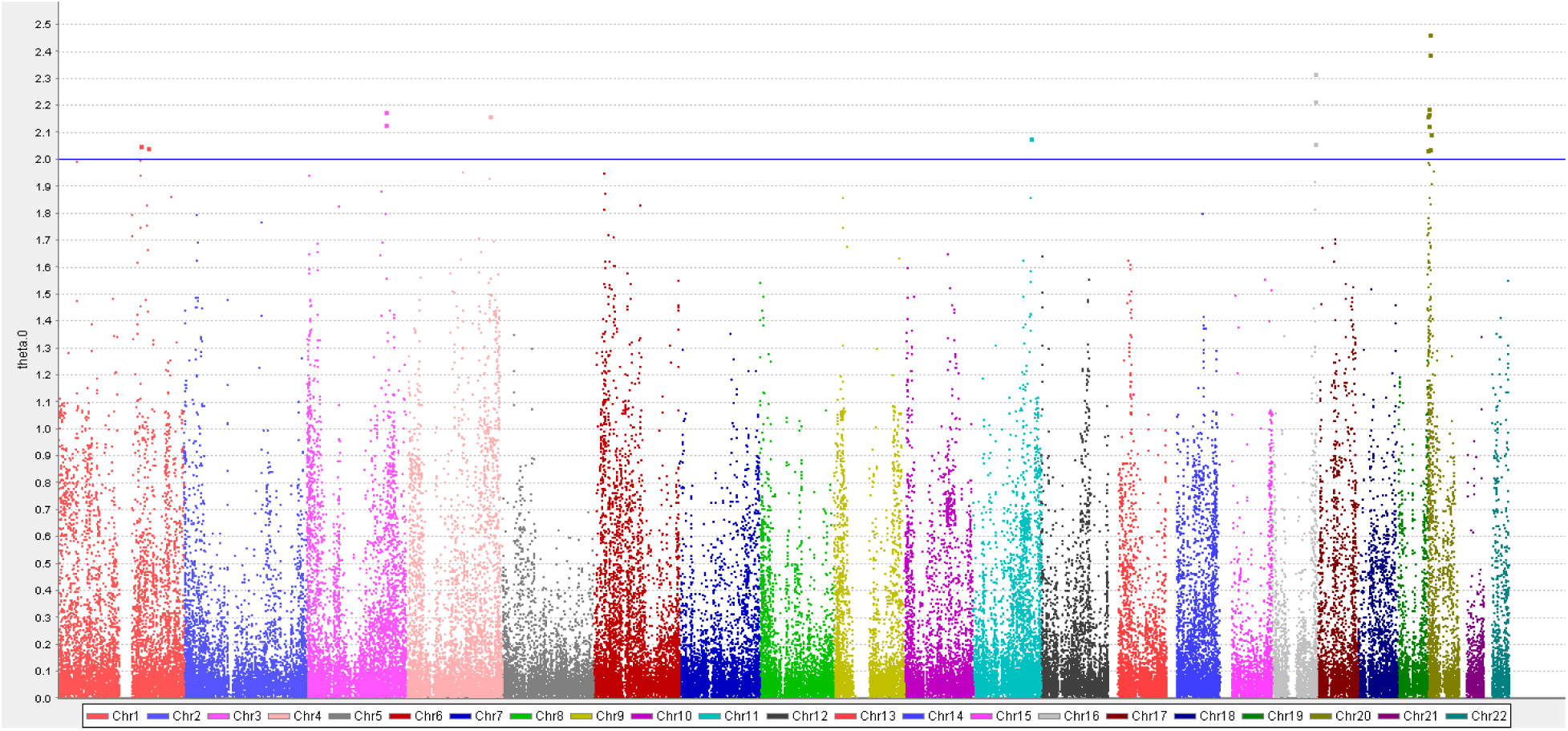
Genome-wide Fastlink two-point LOD scores for affecteds-only model at theta=0. The top three highest LOD scores are all derived from 20p SNPs. Two SNPs (rs8123671 and rs6116410) with a highest LOD score of 2.463 are only <0.6Mb away from TMEM230.

**Fig. 3.**
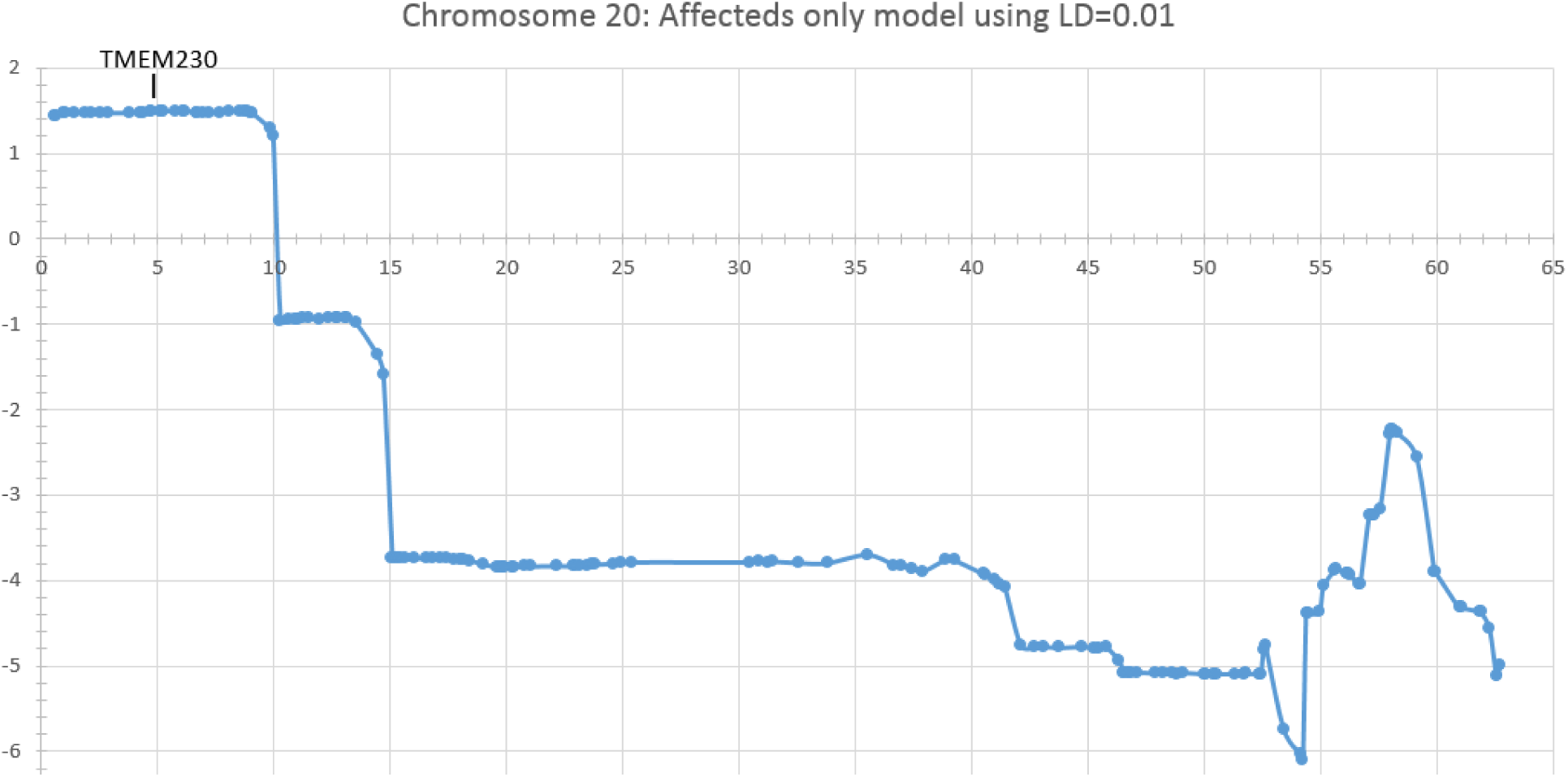
Multipoint LOD scores of affecteds-only model on chromosome 20. A cluster of positive LOD scores of 1.50 was observed in the region from the 0-10Mb region of the short arm of chromosome 20, which almost exactly overlaps with the PD locus (0-10.7Mb, including TMEM230 at 5.1Mb) that we previously defined using microsatellite markers.

**Fig. 4.**
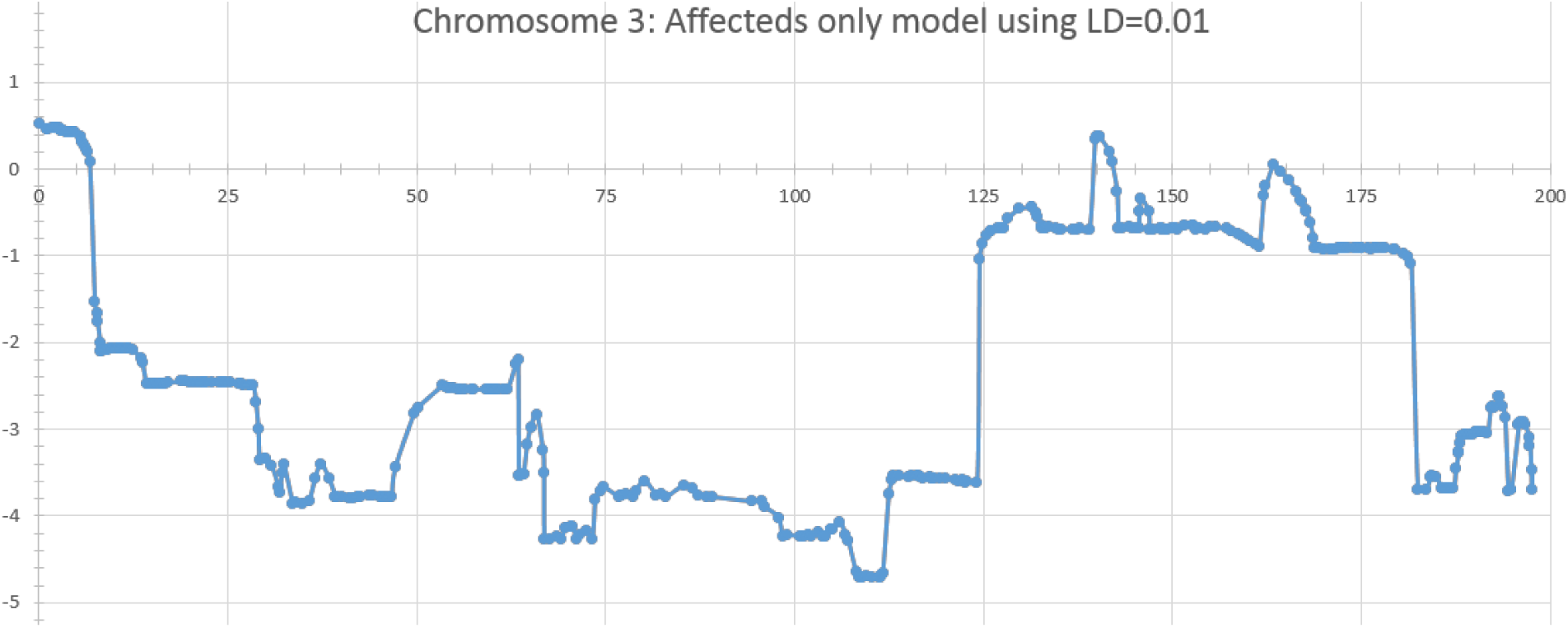
Multipoint LOD scores of affecteds-only model on chromosome 3. The *DNAJC13* locus (132.5Mb) on chromosome 3 yielded negative LOD scores (- 0.4).

**Table 2.** Top 10 genome-wide two-point LOD scores for affected-only model at theta=0

| CHR | BP | SNP | theta=0 | theta=0.05 | theta=0.1 | theta=0.15 | theta=0.2 | theta=0.3 | theta=0.4 |
| --- | --- | --- | --- | --- | --- | --- | --- | --- | --- |
| 20 | 4516796 | rs8123671 | 2.463 | 2.221 | 1.967 | 1.702 | 1.423 | 0.829 | 0.252 |
| 20 | 4522535 | rs6116410 | 2.463 | 2.221 | 1.967 | 1.702 | 1.423 | 0.829 | 0.252 |
| 20 | 4100615 | rs16989226 | 2.391 | 2.152 | 1.901 | 1.639 | 1.365 | 0.785 | 0.234 |
| 16 | 84961276 | rs12928583 | 2.318 | 2.061 | 1.795 | 1.519 | 1.236 | 0.664 | 0.183 |
| 16 | 83983755 | rs2245008 | 2.215 | 1.955 | 1.686 | 1.409 | 1.126 | 0.568 | 0.138 |
| 20 | 1543409 | rs2263664 | 2.188 | 1.978 | 1.752 | 1.512 | 1.256 | 0.71 | 0.203 |
| 3 | 157240416 | rs9855702 | 2.176 | 1.929 | 1.678 | 1.425 | 1.172 | 0.672 | 0.22 |
| 20 | 1913481 | rs6105944 | 2.169 | 1.97 | 1.748 | 1.505 | 1.243 | 0.681 | 0.179 |
| 4 | 165422101 | rs1505392 | 2.163 | 1.918 | 1.669 | 1.419 | 1.169 | 0.675 | 0.224 |
| 20 | 593308 | rs6139718 | 2.16 | 1.952 | 1.73 | 1.493 | 1.241 | 0.701 | 0.199 |

Note. Among top 10 genome-wide two-point LOD scores for 13 affected-only model (SNP genotyping data from II-4 and III-26 did not pass quality control), six were derived from SNPs on chromosome 20p. Importantly, all top three LOD scores were derived from SNPs on our previously assigned PD locus (0-10.7Mb) of chromosome 20p. A highest LOD score of 2.463 was shown at rs8123671 and rs6116410, which are only less than 0.6Mb away from TMEM230-R141L mutation. Note: a SNP (rs9855702) from chromosome 3q showed LOD score of over 2.176. But this SNP (rs9855702, at the 157.2 Mb position) does not represent *DNAJC13* locus (at the 132.5Mb position), and is 24.7Mb away from *DNAJC13*.

Taken together, our whole-genome linkage analysis with ∼300 microsatellite markers and ∼2 million SNPs revealed that chromosome 20p remains to be the most likely PD locus for the Mennonite PD family; re-analysis of the whole-exome sequence data suggested TMEM230-R141L to be the best explanation of etiology for PD in the Mennonite PD family; identification of additional mutations in multiple PD families in different populations **alone** is **sufficient** genetic evidence to establish the pathogenicity of *TMEM230* mutations to PD. Based on these multiple lines of robust genetic evidence, we are confident to conclude that *TMEM230* is a new PD-causing gene. Further studies of *TMEM230* should provide important mechanistic insights into understanding the vesicle/endosome trafficking dysfunction in the pathogenesis of PD, with implications in therapeutic development for PD in the future.

We have updated and revised original Supplementary Fig. 1, which is attached at the end of this response. Our point-to-point responses to the comments from Dr. Farrer et al are listed below.

Han-Xiang Deng, MD, Ph.D. & Teepu Siddique, MD.

Neurogenetics Laboratory, Ken and Ruth Davee Department of Neurology

Northwestern University Feinberg School of Medicine

Tarry Building 13-715

303 E. Chicago Ave

Chicago, IL 60611, USA

Point-to-point responses to Dr. Farrer’s comments

**Comment-1** Deng et al. report the discovery of TMEM230 c.422G>T (p.Arg141Leu) mutation as a cause of late-onset, autosomal dominant Parkinson’s disease (PD) 1 in the same pedigree in which we previously assigned DNAJC13 c.2564A>G (p.Asn855Ser) as pathogenic 2. The chromosome 20pter-p12 locus was discovered by short tandem-repeat (STR) genotyping and linkage analysis, with subsequent exome sequencing in four affected (II-4, III-1, III-20 and III-26) and one unaffected family member. Deng et al. state the rationale for their re-analysis was the inconsistency of genotype-phenotype correlations for DNAJC13 c.2564A>G (p.Asn855Ser), this mutation being absent in three affected family members (II-1, III-1 and III-23). However, two of these suffer atypical parkinsonism; II-1 had clinical and pathologically-proven progressive supranuclear palsy, not Lewy body PD, whereas his son developed symptoms more than two decades earlier than the mean age at onset in the family. Based on haplotype and Sanger sequence analysis the authors’ claim TMEM230 c.422G>T fully co-segregates with disease. Further discussion of these authors’ claims is necessary.

**Response 1.** First of all, we would like to make the facts clear that we did not do the genetic re-analysis of the large Mennonite PD family because we found the inconsistency of genotype-phenotype correlations for *DNAJC13* c.2564A>G (p.Asn855Ser) reported by Dr. Farrer’s group. We began collection of blood samples from the members of this family from 1992. Collection of blood samples from 65 members of this family was completed by 1996, sixteen years before Dr. Farrer’s group started to study this family. We were first supported by the American Parkinson’s Disease Association in 1998 to do linkage analysis for mapping the PD locus to a specific chromosome region, which represented the first step for disease gene identification in the classical genetic studies before exome sequencing approach became available. We mapped a new PD locus to the short arm of chromosome (20pter-p12, 0-10.7Mb region) and started to sequence individual genes in the 10.7Mb minimum candidate region using classical Sanger sequencing strategy. When exome sequencing became available, our efforts were supported by the NIH to identify the new PD gene in this family in 2011 (NS074366, Identification of a Novel Gene for Parkinson Disease). We identified TMEM230-R141L mutation in this family in early 2012. After that, we spent nearly five years to find additional and independent genetic evidence for *TMEM230* in PD, and to characterize *TMEM230*, as shown in our paper ^1^.

Indeed, when Dr. Farrer’s group reported DNAJC13-N855S mutation in this family in 2014 ^2^, we noticed that they identified a different mutation, other than TMEM230-R141L, in this family. But the DNAJC13-N855S was apparently inconsistent with genotype-phenotype correlations, as shown to be absent in three affected family members (II-1, III-1 and III-23). Dr. Farrer’s group explained these inconsistencies by two phenocopies (III-1 and III-23) and a different disease entity (PSP) from PD (II-1). They assumed III-1 (the affected son of the PSP case) to be a phenocopy, mainly because he had a relatively early onset (49 years) than the others. In fact, the age at onset of III-1 is one year older than another affected individual II-16, who had disease onset at 48 years. Moreover, III-1 have typical clinical symptoms of PD. He is now in Hoehn & Yahr stage 5 (advanced disease stage). They also assumed III-23 to be another phenocopy, although he has all cardinal signs of PD with the age of disease onset of 68 years.

Dr. Farrer also argued that the PSP (II-1) case should be excluded, based on the assumption that PD and PSP have different etiology. Indeed, II-1 developed clinical Parkinsonism features. Pathological examination revealed depigmentation and severe degeneration of the substantia nigra, together with tau pathology that was compatible with PSP. Since the PSP case (II-1) died and pathological study was performed in early 1996, before alpha-synuclein pathology was reported in PD in 1997 ^3^, as such the brain of II-1 was not examined for alpha-synuclein pathology. However, the son of the PSP patient, that is III-1, developed clinically typical, levodopa-responsive PD at age of 49 years.

We currently don’t know whether PD and PSP share an etiology in some cases, or PD genetic defects increase the risk to PSP. However, increasing evidence has merged to support this possibility. It has been shown that over 9% of the clino-pathologically confirmed PD cases had co-existing PSP pathology ^4^. Moreover, recent data from genetic and clino-pathological studies support that PD and PSP may be etiologically linked in some cases with well-established PD-causing mutations in *LRRK2*, *SNCA* and *RAB39B*. In these cases, tau pathology might co-exist with alpha-synuclein pathology, or only tau pathology was observed without alpha-synuclein pathology, as detailed below.

1. Tau pathology was found in the brains of patients with SNCA-A53T in the Contursi kindred, and Iowa kindred with SNCA triplication ^5-7^. The tau pathology might not be attributed to old age, because the patient with SNCA-A53T in the Contursi kindred was only 49 years old at autopsy ^5^.
2. In the original PD family D (Western Nebraska kindred) with LRRK2-R1441C mutation, four patients with this mutation were autopsied. Two cases showed Lewy body pathology. The third case had “nonspecific” substantia nigra degeneration with ubiquitin-positive neuronal inclusions. However, the fourth case showed ***only*** tau pathology similar to PSP, and no Lewy body pathology was found ^8,9^. Based on this unusual finding, the authors proposed that LRRK2 functional defects may also be related to other neurodegenerative disorders, such as PSP. In fact, this is similar to what we observed in the Mennonite PD family, one out of four autopsy samples showed PSP pathology. The other three showed Lewy body pathology.
3. An LRRK2-R1441H mutation was found in a family with multiple PD cases. One of the patients had clinically typical PD for 8 years, then transited to PSP during 11 years of the disease course ^10^.
4. In a family with multiple PD cases with LRRK2-G2019S mutation, one patient had clinical PD without ophthalmoplegia. At autopsy, however, tau pathology similar to PSP was found. No Lewy body pathology was observed ^11,12^. **The authors of this paper, including Dr. Farrer, also proposed that Lewy body pathology and tau pathology “may share the same primary cause” ^12^**.
5. Loss of function mutations in *RAB39B* have been shown to cause X-linked intellectual disability and early-onset PD with alpha-synuclein pathology ^13^. In the autopsy sample from a patient with full deletion of *RAB39B*, Wilson et al found both alpha-synuclein pathology and tau pathology ^13^. Again, the tau pathology might not be attributed to old age, because the patient died at 48 years old from positional asphysia^13^. Taken together, although the total number of the autopsy cases with well-established pathogenic mutations in *SNCA*, *LRRK2* or *RAB39B* is still very limited (∼50 cases) in the literature to date, tau pathology similar to PSP has been found in a significant fraction of these cases (∼10%). **This raises the possibility that PD and PSP may indeed share potential etiology, or the PD-causing mutations increase PSP risk, at least in some cases**. Additional support of this possibility includes: (I) We have two autopsies on two real brothers, clinically and pathologically one had Parkinson’s disease and the other PSP, although the genetic cause remains unknown; (II) On reading our paper, Dr. Stanley Fahn, a renowned PD expert, informed us on a family with PD and PSP in a parent–child pair; (III) Mutations in *MAPT* cause a subtype of parkinsonism linked to FTDP-17 ^14^; (IV) Tau-positive pathology without Lewy body has been observed in the substantia nigra of multiple cases with sporadic, but clinically diagnosed PD ^15^. Genetic pleomorphism is not an uncommon observance. For example, mutations in *VCP* give rise to inclusion body myopathy with Paget disease of the bone and frontotemporal dementia ^16^, amyotrophic lateral sclerosis ^17^, and Charcot-Marie-Tooth Disease Type 2Y ^18^; *LMNA* mutations give rise to 12 distinct disease phenotypes involving different organs and systems with autosomal dominant and/or recessive patterns ^19^.

It is important to note that Farrer’s group reported in their paper that they sequenced the entire coding region of 115 genes known to be implicated in neurological disorders in their three assumed “phenocopy” cases (II-1, III-1 and III-23) using a targeted capture panel ^2^. Despite this effort, none of the identified variants could account for disease in these three patients ^2^. These data clearly indicate that a new genetic variant, other than DNAJC13-N855S (or any variants in these 115 genes), is responsible for PD in this family. TMEM230-R141L mutation provides a parsimonious explanation for PD in this family, and was found in all three affected patients (II-1, III-1 and III-23). In addition, even if PSP is not in the spectrum of genetic pleomorphism associated with canonical PD genes, then II-I can still be considered an obligate career, having passed on the PD trait to his offspring, as shown in the affected PD case (III-1).

**Comment-2.** Our prior exome analysis of III-15 identified a DNAJC13 c.2564A>G (p.Asn855Ser) mutation but not TMEM230 c.422G>T (p.Arg141Leu) (Figure 1a). The exons of both genes had >30x sequence coverage and nucleotide sequences at both sites were confirmed by Sanger sequencing. In addition, our chromosome 20 STR marker genotyping, while consistent with parental/sibling haplotypes and Mendelian inheritance (Figure 1a) shows the genotyping presented by Deng et al. is erroneous for two individuals; III-14 at age 75 years is an unaffected carrier of TMEM230 c.422G>T (p.Arg141Leu) whereas III-15 is wild-type (Figure 1a, compare with Supplementary Fig. 1). Many of the samples we have examined were obtained twice including III-15; blood was collected prior to 2010 and at a family reunion organized in 2013, and these samples gave identical results. Nevertheless, phenocopies in large pedigrees with multi-incident, monogenic late-onset parkinsonism are to be expected.

**Response-2.** As pointed out by Dr. Farrer, we have updated the affection status based on Dr Rajput’s most recent input. We incorrectly assigned the disease status to III-15 in the previous Supplementary Fig.1. Indeed, the individual III-14, but not III-15, was affected. The clinical information in our records dated back to 2009, and has now been updated (Supplementary Fig. 1). This new information, in essence, does not change the status of TMEM 230-R14L mutation as causative in the family.

The clinical symptoms of the affected III-14, the only phenocopy for TMEM 230-R141L, appear to be very unusual. Sixteen years after he showed initial left thumb tremor at 59 years old, clinical diagnosis of parkinsonism could not be made by Dr. Ali Rajput. We obtained the last updated clinical information when he was evaluated in 2009 as uncertain status. In 2014, when Vilarino-Guell et al ^2^ reported DNAJC13-N855S in this family with newly updated clinical information, they had more recent clinical information.

Based on Dr. Rajput’s recent evaluation, III-14 is now 82 years old. After 23 years since he started to show very mild and questionable signs when he was 59 years old, he is still not on L-DOPA. This is the only affected case without TMEM230-R141L mutation in this family. Therefore, for TMEM230-R141L mutation, a single phenocopy event is needed to explain the phenotype of individual III-14.

As a late-onset and relatively common disease, PD has been reported to have a high phenocopy rate ^20^. It has been shown that in the families with genetically defined forms of PD, phenocopy may occur at the remarkable rate of 14.4% in well-described pedigrees, and in 5.0% of all affected individuals in pedigrees ^20^.

Indeed, the pathogenicity of TMEM230-R141 also assumes a single phenocopy (II-14) to explain PD in the Mennonite PD family. However, DNAJC13-N855S needs to assume three phenocopies (II-1, III-1 and III-23) to explain the same family. Thus the phenocopy rate for TMEM230-R141L mutation is 6.7% (1/15), which is compatible with 5% of all affected individuals in the pedigrees shown by a previous study ^20^. However, the phenocopy rate will be 20% (3/15) if assuming DNAJC13-N855S mutation is disease-causing. Even if the individual II-1 (PSP case) is not taken into account, DNAJC13-N855S needs to assume two phenocopies to explain PD this family.

As Dr. Farrer realized “Nevertheless, phenocopies in large pedigrees with multi-incident, monogenic late-onset Parkinsonism are to be expected”. However, it appears to be the only genetic evidence that Dr. Farrer used to oppose *TMEM230* as the new PD gene. If Dr. Farrer recognizes phenocopy in PD in this Mennonite family, he should accept the gene with fewer phenocopy (a single phenocopy for TMEM230, II-14), rather than the gene with more phenocopies (at least two phenocopies for DNAJC13, II-23 and III-1, even if II-1 is not taken into account) as the disease-causing gene in this family. Using the precept of Occam’s razor, **among competing hypotheses, the one with the fewest assumptions should be selected**. Among TMEM230-R141L and DNAJC13-N855S, TMEM230-R141L mutation is apparently the more parsimonious explanation for the occurrence of PD in this large family.

Dr. Farrer’s group performed exome sequencing using three affected individuals, including III-14 ^2^. Individual III-14 is the only phenocopy case without TMEM230-R141L among all affected members in this family. **This appears to be the reason why Dr. Farrer’s group missed the TMEM230-R141L as the pathogenic mutation in this family**.

Taken together, all these data support that TMEM230-L141L, rather than DNAJC13-N855S (or ZBTB38-R264Q), is the most likely disease-causing mutation in this large family. The other genetic variants are even more unlikely, because they are absent in five or more affected members in this family (Table 1 and detailed in RESPONSE-3).

**Comment-3.** Deng *et al.* state *DNAJC13* c.2564A>G is present in an asymptomatic individual (II-9), who died at 87 years, and they excluded variants present in this exome and shared by the intersection of four affected exomes. We were not aware of a DNA sample from this subject but as reduced penetrance is a feature of monogenic parkinsonism. We appreciate ‘disease-discordant’ analysis is best avoided. Indeed the authors’ observe *TMEM230* c.422G>T (p.Arg141Leu) in 20/23 subjects including 12 unaffected carriers in generation III. The number of carriers is consistent albeit greater than one might expect for a dominant pattern of disease inheritance. However, the authors’ fail to disclose that many *TMEM230* c.422G>T carriers are advanced in age with 7 individuals beyond the mean age of symptom onset (mean age of unaffected carriers= 69.6±8.2, range 57-84 years; Table). Given these issues segregation of the *TMEM230* c.422G>T (p.Arg141Leu) with parkinsonism is arguably less compelling than for *DNAJC13* c.2564A>G (p.Asn855Ser). Two-point linkage analysis of the chromosome 3q21.3-q22.2 disease-segregating haplotype (also harboring and *ZBTB38* c.791G>A (p.Arg264Gln) variants, 9Mb apart) yielded a LOD=4.79 (Ɵ=0.05) which increased to 5.29 (Ɵ=0) assuming a 10% phenocopy rate. Hence, other considerations for the pathogenicity of these variants must be carefully reviewed.

**Response-3.** Individual II-9 was clinically not affected until she died at 87 years old. We obtained her blood sample in 1994, one year before her death. This individual had DNAJC13-N855S mutation, but did not have TMEM230-R141L mutation. Dr. Farrer raised the concern that the DNA samples of an unaffected (II-9) and a PD patient (III-1, son of the PSP patient) were used for exome sequencing and filtering. This is best avoided. To address Dr. Farrer’s concern, we re-analyzed our whole-exome sequencing data. We selected two individuals: II-4, who developed Lewy body-confirmed PD; and III-26, who developed typical clinical PD, and his father (II-14) had Lewy body-confirmed PD. We identified 12 additional rare variants shared by these two affected individuals. We performed Sanger sequencing study in 13 affected individuals (whose DNA samples are available) for co-segregation analysis (Table 1). We found that 10 variants were not present in at least five affected individuals. Therefore, we excluded these 10 variants as PD-causing in this family.

TMEM230-R141L mutation was detected in all affected individuals, except for III-14. DNAJC13-N855S and ZBTB38-R264Q were not present in three affected individuals, including two known PD cases (III-1 and III-23) and the clinical PD/pathological PSP (II-1) case. We believe that absence of DNAJC13-N855S and ZBTB38-R264Q in three affected members in two different branches (III-23 in one branch, and II-1 and III-1 in the other branch) in this family basically excludes the DNAJC13-N855S and ZBTB38-R264Q as pathogenic variants in this family (Table 1).

Dr. Farrer raised a concern regarding the 12 unaffected members with TMEM230 mutation in the third generation. Most members with TMEM230-R141L mutation in the third generation are still in their 50s to 70s. It is currently unknown whether they will develop PD later in their lives. The affected status of the second generation provides meaningful information in terms of the disease onset in this family, because all eight members in the second generation have passed away at the ages from 78 to 94 years old. All six members with TMEM230-R141L mutation developed either PD (II-4, II-6, II-8, II-11 and II-14) or PSP (II-1). The age at onset of PD ranges from 68 to 85 years (74.3 ± 8.6, mean ± sd). Two members (II-9 and II-12) died without PD. II-12 did not carry either TMEM230-R141L or DNAJC13-N855S mutation, died at 78 years old. However, II-9 carried DNAJC13-N855S, but not TMEM230-R141L mutation, died at 87 years old without PD.

In the third generation, there are 42 family members (excluding spouses). To provide clarity to Supplementary Fig. 1, we added additional information from 28 members, with 19 carriers of the affected TMEM230 haplotype. Most members without the affected haplotype were not included. The major reason why there are more carriers of the TMEM230-R141L than the carriers of the DNAJC-N855S mutation is because six children (III-1 to III-6) of affected individual II-1 carry the TMEM230-R141L mutation. Among these six, one has already developed clinical PD (III-1).

Altogether, there are 50 family members (excluding spouses) in the second and third generations in this family. Nineteen are asymptomatic TMEM230-R141L carriers, and ten are asymptomatic DNAJC13-N855S carriers. In fact, seven out of the ten DNAJC13-N855S asymptomatic carriers also carry the TMEM230-R141L (7/10). Therefore, these seven carriers are not able to provide favorable information for either of these two mutations. Excluding these seven carriers with both mutations, nine are TMEM230-R141L-alone carriers (59-82 years old), with only three being ≥74.3 years old, which is the average age of onset of the PD cases in the second generation. These three TMEM230-R141L carriers are 75, 81 and 82 years old, respectively (the rate of unaffected cases with the age of ≥74.3 years old: 3/9=33.3%). Although there are only three DNAJC13-N855S-alone carriers (66, 80 and 87 years old, respectively), two of them are older than 74.3 years (80 and 87 years old, respectively, 2/3=66.7%). Moreover, the oldest asymptomatic TMEM230-R141L carrier (82 years old) is younger than the oldest asymptomatic DNAJC13-N855S carrier (87 years old). We understand that the number of cases with the age over the average age at onset (74.3 years in the second generation) in the entire family is still small, but based on the limited data, the rate (33.3% for TMEM230-R141L vs 66.7% for DNAJC13-N855S) of the asymptomatic cases over 74.3 years does not favor the pathogenicity of DNAJC13-N855S in this family either.

In terms of the LOD scores, we calculated the LOD scores by unbiased whole genome linkage study using common microsatellite markers before the disease gene was identified ^1^; whereas, Vilarino-Guell et al calculated the LOD score using the extremely rare variants (DNAJC13-N855S and ZBTB38-R264Q) after the variants had been identified ^2^. It is known that LOD scores will be greatly different if the parameters used for calculation are different. The detailed parameters for calculating the LOD scores were not presented by Vilarino-Guell et al ^2^. However, based on the facts that there were more “phenocopies” for DNAJC13-N855S, and there are more affected individuals with TMEM230-R141L than those with DNAJC13-N855S, the LOD score of the DNAJC13-N855S should **never** be higher than the LOD score of TMEM230-R141L in any circumstances, if the same parameters were used for LOD score calculation.

To comprehensively address Dr. Farrer’s comment, in collaboration with Dr Margaret Pericak-Vance, we further performed the whole-genome SNPs genotyping using MegaEx chip. The MegaEx chip contains ∼2 million SNPs. We analyzed 44 sampled individuals in this PD family (#9853). The SNP data from three individuals, including two affected (II-4 and III-26) and one unaffected (III-7), did not pass quality control due to low call rate. The data from all the other 41 individuals were further processed. In the overall dataset, a total of 1,933,857 SNPs had a genotyping rate of >=97%. After selection of polymorphic SNPs and exclusion of very rare SNPs in the family, a final set of 198,493 SNPs were used for analysis.

Mendelian inconsistencies were evaluated and cleaned using PLATO. No problems were indicated with the pedigree structure. We used a set of 99 unrelated Caucasian 1000-Genomes samples for allele frequency estimation and to determine linkage disequilibrium (LD). All SNPs were evaluated for 2-point results using Fastlink. LD pruning was used to select SNPs for multipoint analysis, with an r2 threshold varying from 0.01 to 0.16 for multipoint analysis on chromosomes 3 and 20, resulting in 7,671 SNPs selected for chromosome 3 and 3,194 SNPs selected for chromosome 20. A disease allele frequency of 0.001 was used. Both an affecteds-only and a model with 6 age-based liability classes were run for all analyses. A phenocopy rate of 0.05 was also incorporated into the model. Two-point LOD scores were evaluated at Ɵ=0 and run for both the age-dependent penetrance model and affecteds-only.

In the age-dependent penetrance model, LOD scores of over 2.0 were observed at 11 clusters of the entire genome, including those on chromosome 20p, the region containing *TMEM230* (Fig. 1). The positive cluster on chromosome 20p is located within the PD locus (10.7Mb minimum candidate region from 20pter-p12), a region previously assigned by us using microsatellite markers. However, the positive cluster on 3q is over 25Mb away, distal to *DNAJC13* (Fig. 1). In the affecteds-only model, among the top 10 two-point LOD scores, six were derived from SNPs on chromosome 20p (Fig. 2 and Table 2). Notably, the top three LOD scores were all derived from SNPs on chromosome 20p. A highest LOD score of 2.463 was observed at rs8123671 and rs6116410, which are only less than 0.6Mb away from TMEM230-R141L mutation (Table 2). If fact, the LOD scores would be expected to be even higher if SNP data from two affected individuals with TMEM230-R141L mutation (II-4 and III-26) had been informative. Although an SNP (rs9855702) from chromosome 3q showed LOD score over 2.176 (Table 2), again, this SNP is 25Mb away from DNAJC13-N855S.

To further address the linkage of PD to *TMEM230* locus on chromosome 20 or to *DNAJC13* locus on chromosome 3, we performed Merlin Multipoint analysis using the affecteds-only model. A cluster of positive LOD scores of 1.5 was observed in the region from 0-10Mb of the short arm of chromosome 20 (Fig. 3), which almost exactly overlaps with the PD locus (0-10.7Mb) that we previously defined using microsatellite markers ^1^. In contrast, the *DNAJC13* locus (132.5Mb) yielded negative LOD scores of −0.4 (Fig. 4).

Overall, with the same parameters for calculation of the genome-wide SNPs, the chromosome 20p (0-10Mb, *TMEM230* locus) yielded consistent positive two-point LOD scores over 1.5, with a highest two-point LOD score of over 2.46 in the affected-only model shown at rs8123671 and rs6116410, which are only less than 0.6Mb away from TMEM230-R141L mutation. Although high two-point LOD scores were also observed in a few regions of the other chromosomes in the age-dependent penetrance model, these scores were greatly diminished in the affected-only model. In contrast, the SNPs around *DNAJC13* locus yield consistently much lower LOD scores in any models. Moreover, the *DNAJC13* locus (132.5Mb) yielded negative multipoint LOD scores of −0.4 in the affecteds-only model.

In summary, our new data from whole-genome SNPs genotyping using MegaEx chip demonstrate the strongest linkage of PD to chromosome 20p (0-10Mb), including *TMEM230* in the Mennonite PD family. These new data are consistent with our previous linkage data using whole genome microsatellite markers. One phenocopy for TMEM230-R141L vs three phenocopies for DNAJC13-N855S clearly reflects the obvious difference (Table 1). Therefore, we believe there is no clear evidence at the linkage or at whole-exome sequencing level to support Dr. Farrer’s claims that the *DNAJC13* locus has “higher” LOD scores than *TMEM230* locus, and “genetic support for *DNAJC13* remains ostensibly greater.” The best genetic explanation for PD in this Mennonite family remains to be the TMEM230-R141L mutation at the 20p locus.

**Comment-4.** The *DNAJC13* c.2564A>G mutation is not observed in public databases (ExAC 0.3.1 release Mar 14^th^ 2016) and the amino acid codon it encodes, p.Asn855, is evolutionarily conserved across species, suggesting it is important to protein function. In contrast, while *TMEM230* c.422G>T is novel, its codon p.Arg141 is not evolutionarily conserved in humans. This is an important caveat. TMEM230 p.Arg141Gln and p.Arg141Trp substitutions occur with appreciable frequency; in Caucasians their combined minor allele frequency (MAF) > 5e-5 and in South Asia TMEM p.Arg141Trp MAF=0.00024 alone (ExAC 0.3.1 release Mar 14th 2016). CADDv135, which provides a composite prediction of damage to protein structure, suggests these substitutions are comparable (CADDv13 scores for DNAJC13 p.Asn855Ser =23.6, and for TMEM230 p.Arg141Leu, p.Arg141Gln and p.Arg141Trp = 27.6, 23.8 and 28.6 respectively). While the MAF for TMEM p.Arg141 substitutions is more than one might envisage for a rare cause of parkinsonism the clinical phenotype of these carriers is unavailable and *in-silico* predictions cannot be assumed causal of disease.

**Response-4.** Evolutionary conservation should be considered between species during evolution. The amino acid at R141 is invariable in all the known species during evolution. Indeed, R141Q variant is present in two and R141W variant in five individuals among over 60,000 individuals in the ExAC database. These two variants are present in South Asian and Latino, not in European samples. They are very rare, and their pathogenicity remains currently unknown. However, they are different variants from R141L. Different variants at the same amino acid position are expected to have variable functional consequences. As shown in Table 3, two widely used pathogenicity programs consistently predicted R141L to be damaging with high confidence, while the other two variants were either inconsistent for pathogenicity (R141W) or consistently benign (R141Q) (Table 3). Notably, Vilarino-Guell et al identified DNAJC13-R2115L mutation in a small family (TU1) with two affected ^2^. In fact, at the same codon, another variant, DNAJC13-R2115Q, is also present in five individuals in the ExAC database.

**Table 3.**
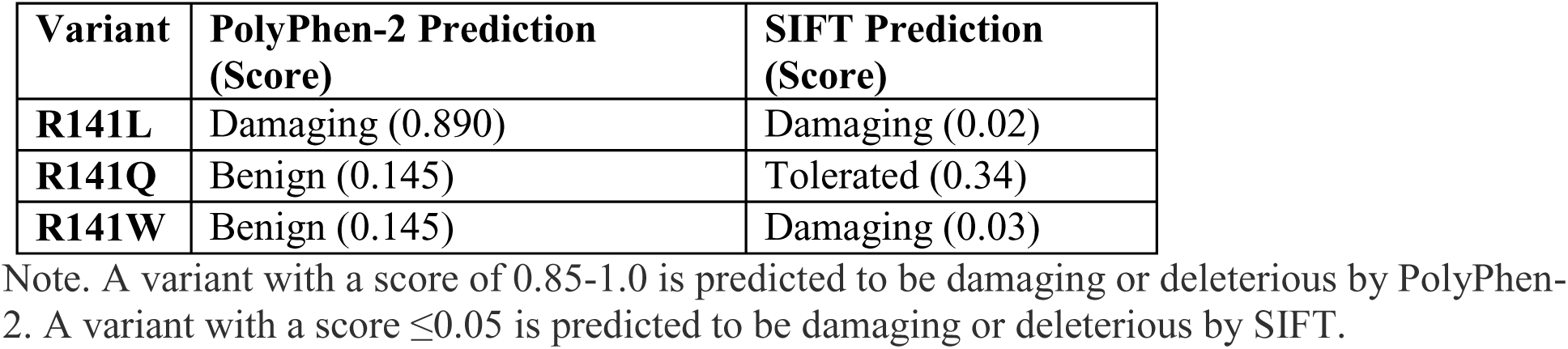
Pathogenicity prediction of variants at TMEM230-R141

| **Variant** | **PolyPhen-2 Prediction**<br>**(Score)** | **SIFT Prediction**<br>**(Score)** |
| --- | --- | --- |
| **R141L** | Damaging (0.890) | Damaging (0.02) |
| **R141Q** | Benign (0.145) | Tolerated (0.34) |
| **R141W** | Benign (0.145) | Damaging (0.03) |

Note. A variant with a score of 0.85-1.0 is predicted to be damaging or deleterious by PolyPhen-2. A variant with a score $\leq 0.05$ is predicted to be damaging or deleterious by SIFT.

**Comment-5.** For Nature Genetics ‘proof of pathogenicity’ requires additional mutations in the same gene segregate with disease in independent families. In our original study screening of *DNAJC13* c.2564A>G (p.Asn855Ser) identified five more affected carriers but no controls; three of the positive cases have a family history of parkinsonism and in two small pedigrees co-segregation with PD was evident. All DNAJC13 p.Asn855Ser carriers are of Dutch-German-Russian Mennonite origin and share a common haplotype.

**Response-5.** DNAJC13-N855S mutation was detected in the original Mennonite family, but with three affected individuals lacking this mutation. Genetic study of a large cohort of 2,928 PD cases by Vilarino-Guell et al revealed that this mutation was present in four single PD cases. No co-segregation of this mutation with PD in any families could be shown^2^. The same research group subsequently performed sequencing and genotyping analysis with a large cohort of 1,964 parkinsonism cases from Canada and Norway, and they only found a single case with the DNAJC13-N855S mutation ^21^. There is no evidence of DNAJC13-N855S co-segregation with PD in any families, except for the incomplete segregation in the original Mennonite family. Dr. Farrer mentioned “In our original study screening of *DNAJC13* c.2564A>G (p.Asn855Ser) identified five more affected carriers but no controls; three of the positive cases have a family history of parkinsonism and in two small pedigrees co-segregation with PD was evident.” But in fact, in family BC1, DNAJC13-N855S was present in one PD case only; the other is a dementia case without PD; and in TU1 family, another variant, DNAJC13-R2115L (but not DNAJC13-N855S) was detected in the affected mother and son. But the son with DNAJC13-R2115L variant developed disease very early, at 24 years old; whereas, his sister carrying the same variant was unaffected by age 46 years old. Because the pathogenicity of DNAJC13-R2115L to PD could not be established by such a single and small family, this variant may not provide meaningful supporting evidence for the pathogenicity of DNAJC13-N855S to PD either.

Altogether, the DNAJC13-N855S was found in five out of a total of 4,892 PD cases tested by Dr. Farrer’s group. This variant is only present in the cases with Mennonite heritage (with the same haplotype). Therefore, it is highly likely that the DNAJC13-N855S is a private variant specific to the Mennonite ethnic population, but not pathogenic for PD in this family. Indeed, as Dr. Farrer and his co-authors noted, “additional proof for pathogenicity is needed” for DNAJC13-N855S mutation ^2^. Also, in Dr. Farrer’s 2013 review of genetics of Parkinson disease, he and his co-author realized that the DNAJC13-N855S mutation “could reflect a unique benign variant originating from a common founder that does not influence disease” ^22^. This notion is supported by worldwide genetic studies of PD to date.

1. Ross et al sequenced the entire coding exons in a French-Canadian/French cohort of 528 PD patients, but did not find any pathogenic mutations, and none of the variants identified in their study were independently associated with PD ^23^.
2. Lorenzo-Betancor et al sequenced DNAJC13-N855S variant in a very large Caucasian series consisting of 1,938 cases with clinical PD and 838 with pathologically diagnosed Lewy body disease, they did not identify DNAJC13-N855S variant in any of these 2,776 cases ^24^;
3. Foo et al sequenced all coding exons of DNAJC13 in 99 Chinese PD cases and subsequently genotyped a large cohort with 711 PD cases. They did not identify any pathogenic mutations, and none of the variants identified in their study showed any association to PD ^25^.

Overall, a total of 4,114 cases have been sequenced or genotyped for variants in *DNAJC13* by other investigators. DNAJC13-N855S has not been identified in any additional PD cases. Moreover, no additional genetic studies have provided supporting evidence for the pathogenicity of *DNAJC13* mutations to PD or association of *DNAJC13* variants with PD in different populations to date ^23-25^.

**In contrast, we identified TMEM230-*184ProGlyext*5 mutation in nine PD patients in seven unrelated Chinese families.** This mutation was shown to co-segregate with PD in two families (Supplementary Figs. 8 and 11) ^1^. We also identified TMEM230-*184Trpext*5 mutation in a family PD patient in North America. Intriguingly, TMEM230-*184ProGlyext*5 and TMEM230-*184Trpext*5 are different mutations in different populations, and both mutations involve the stop codon, adding identical five amino acids (HPPHS) at the C terminus ^1^.

Identification of TMEM230-R141L mutation in the large Mennonite PD family was the original driving force leading us to test TMEM230 mutations in other PD patients. We subsequently identified two different mutations with nearly identical functional impact in the same gene in eight families from two well-separated populations (North America and China) ^1^. **We believe that identification of these two TMEM230 mutations (*184Trpext*5 and *184ProGlyext*5) alone is sufficient genetic evidence to establish the pathogenicity of TMEM230 mutations to PD.**

In addition to our initial report, Baumann et al recently identified TMEM230-R68H mutation in a clinically typical and familial PD case in Germany ^26^. Notably, this PD patient is from a Mennonite family of Ukrainian ancestry without DNAJC13-N855S mutation. Mutations in other known PD-linked or PD-risk genes are negative in this patient ^26^. Most recently, Yang et al screened a Chinese cohort with 355 sporadic PD cases for TMEM230 mutations ^27^. They found TMEM230-A110T mutation in two apparently unrelated PD cases. The amino acid at Thr110 is highly evolutionally conserved. This mutation was neither present in more than 500 Chinese control samples, nor in the ExAC database where more than 60,000 unrelated individual DNA samples have been exome-sequenced ^27^. These new data provide the first genetic evidence that TMEM230 mutations may also be involved in the apparently sporadic PD cases.

**Comment-6.** Our screening of *TMEM230* c.422G>T in 1283 Canadian patients (726 with sporadic PD and 557 with familial parkinsonism), including those of Dutch-German-Russian Mennonite ancestry in our original report, did not identify any carriers. Deng *et al.* go on to identify *TMEM230* c.275A>G (p.Tyr92Cys) and c.551A>G (p.*184Trpext*5) in two patients with early-onset parkinsonism; the former observed in an unaffected maternal carrier and the latter found in a sporadic case. In addition, *TMEM230* c.550_552delTAGinsCCCGGG (p.*184ProGlyext*5) was found in affected probands from 7 of 225 unrelated families of ethnic Han Chinese descent. Nevertheless, while all these mutations are novel, none are shown to segregate with disease.

**Response-6.** As mentioned above, we identified TMEM230-*184ProGlyext*5 mutation in nine PD patients in seven unrelated Chinese families. In fact, **this mutation was shown to co-segregate with PD in two families** (Supplementary Figs. 8 and 11). We also identified TMEM230-*184Trpext*5 mutation in a family PD patient in North America. Intriguingly, both mutations involve the stop codon, adding identical five amino acids (HPPHS) at the C terminus. **We believe that identification of two different mutations with nearly the same functional impact in a total of 10 familial PD patients from 8 families alone is sufficient to establish the pathogenicity of *TMEM230* mutations to PD, regardless the disease mechanisms.**

Also as mentioned above, Baumann et al recently reported TMEM230-R68H mutation in a Mennonite familial PD case without mutations in the other PD genes and DNAJC13 ^26^. Yang et al found TMEM230-A110T mutation in two Chinese apparently sporadic PD cases ^27^.

**Comment-7.** The TMEM230 p.*184ProGlyext*5 discovery is most incongruous given the independent/ancient haplotypes on which it is observed (Supp. Fig 12d) and the 3.1% of familial PD in ethnic Chinese it is stated to explain. In contrast to TMEM230 p.Arg141Leu, the p.*184ProGlyext*5 substitution does not appear to follow a Mendelian pattern of inheritance, although the lack of clinical information for parents and siblings (many of whom must be carriers) offers little insight. We have examined *TMEM230* variability in 256 patients with early-onset, familial and/or atypical parkinsonism; while we observe 4/7 *TMEM230* variants leading to amino acid substitutions (p.Met1Val, p.Arg62His, p.Ile125Met, p.Arg171Cys) all have appreciable frequencies in ExAC (MAF >5e-5) and only p.Arg171Cys is predicted to be damaging to protein structure (MAF=0.003, CADDv13 score=34). We also genotyped *TMEM230* c.550_552delTAGinsCCCGGG in 80 affected probands with familial PD and 280 control subjects of ethnic Chinese descent from Taiwan but found no carriers.

**Response-7.** In our original report, we screened a Chinese cohort with 574 PD cases, including 225 familial index cases and 349 sporadic cases. We identified a single mutation, TMEM230-*184ProGlyext*5 in seven familial index cases from seven apparently unrelated families but not in the sporadic cases. In two families with additional DNA samples available from other PD patients, **we could confirm the co-segregation of disease with this mutation**. To test whether mutations in other known Parkinson’s disease–linked genes, such as *SNCA*, *LRRK2* and *VPS35*, or other parkinsonism-linked genes were present in these Chinese PD patients, we further performed whole-exome sequencing using DNA samples from these nine Chinese PD patients. We did not find new variants in these genes. We performed Sanger sequencing of DNA samples from our 528 in-house Chinese controls, and we did not find this mutation. Moreover, this mutation was not present in the Chinese exome sequencing database, which included over 9,000 neurologically normal Chinese controls, from BGI-Shenzhen, nor in any public databases, including Exome Aggregation Consortium (ExAC) database with a total of 60,706 unrelated individuals. Of the nine PD patients from seven families, five patients from four families were homozygotes for the mutation (Supplementary Figs. 5, 6, 9 and 11). Our haplotype analysis suggested that this mutation either arose independently or is very ancient. The p.*184Trpext*5 and p.*184ProGlyext*5 alterations were identified in different populations (North America and China), but both affect the stop codon, resulting in addition of identical five amino acids (HPPHS) at the C-terminal of the mutant TMEM230. The mutation encoding p.*184ProGlyext*5 accounted for 3.1% (7/255) of Chinese index cases with familial PD in our cohort.

We understand that size of the Chinese familial PD cohort was relatively small (255 familial index cases), and mostly from the middle south/west regions of China. This mutation may not have even distribution in Chinese population. It is not surprising that Dr. Farrer did not detect this mutation in only 80 Chinese familial PD probands from Taiwan. Additional genetic studies from different parts of the Chinese PD cases may provide further information for better understanding the frequency and distribution of *TMEM230* mutations in Chinese PD.

Dr. Farrer raised a concern regarding the inheritance pattern of the mutation encoding TMEM230-*184ProGlyext*5, because “the p.*184ProGlyext*5 substitution does not appear to follow a Mendelian pattern of inheritance”. Indeed, we found that TMEM230-*184ProGlyext*5 in familial PD cases in both heterozygous and homozygous conditions, even in the same families. But this is not unusual. In fact, there are many other examples like this, such as mutations of *SOD1* and *FUS* in ALS. Although mutations in *SOD1* or *FUS* generally cause ALS in a dominant fashion, recessive pattern of inheritance has also been found by some mutations, such as SOD1-D90A^28^, SOD1-delG27/P28 ^29^ and FUS-H517Q^30^. Notably, SOD1-D90A mutation can cause ALS in both dominant (heterozygous condition) and recessive (homozygous condition) fashions, with homozygous mutation appearing to have higher penetrance than heterozygous mutation, likely due to a dose effect ^28,31-34^. Therefore, we suggested in our original report that this may “be due to an increased penetrance of the TMEM230-*184ProGlyext*5 mutation in homozygous condition in later-onset PD due to a dose effect.”

Dr. Farrer also raised a concern regarding whether the occurrence of the TMEM230 p.*184ProGlyext*5 mutation was independent or was very ancient. In fact, it does not appear to be rare for the patients with the same genetic mutation, but with different haplotypes, which is thought to be a sign of the mutation arising independently, or extremely ancient. For examples, Zimprich et al identified the same R1441C mutation in *LRRK2* in two unrelated families, including family D (Western Nebraska) and family 469. Haplotype analysis with five flanking polymorphic markers, including three *LRRK2* intragenic markers, revealed different haplotypes. Similarly, they pointed out that the R1441C mutation either arose independently or extremely ancient ^9^. Another example for independent events for the same genetic mutation is the ALS-linked SOD1-A4V, which represents the most prevalent SOD1 mutation, accounting for approximately half of the ALS patients with *SOD1* mutations in North America ^35^. Our study suggested two separate origins of this mutation, one in Europe and the other in Amerindians, approximately 400-500 years ago ^36^.

**Comment-8.** Deng *et al.* claim TMEM230 mutations impair ‘synaptic vesicle trafficking’. However, assuming the TMEM230 antibodies are specific (which was not demonstrated in the manuscript), TMEM230 localized to large vesicular structures in the cell body. The localization appeared similar for endogenous and overexpressed/tagged protein in Neuro-2a cells and brain tissue (Figures 2 & 3). Similarly, confocal live cell imaging showed the movement of tagged, overexpressed (non-physiological) proteins in neuronal soma (Figure 4). Assessing ‘synaptic vesicle’ transport speed, track length and displacement length is claimed to show some mutant specific differences. Nevertheless, synaptic vesicles by definition reside in presynaptic axon terminals; somatic membrane structures containing vesicular proteins are not synaptic vesicles, which form from membrane intermediates at axonal sites hugely distal from the soma. It is incongruous this biology was not studied in an appropriate, physiological context or at least termed correctly. How images were quantified is unclear; at a minimum transfection levels for each tagged protein must be comparable and should have been demonstrated. The significance of the 1-way ANOVA is also based on comparison to an overexpressed, tagged wild-type protein. The more frequent and presumably non-pathogenic TMEM230 substitutions we have identified might prove to be informative as controls.

**Response-8.** We identified *TMEM230* mutations five years ago. To understand the localization of TMEM230 in neuronal cells and the impacts of its mutations, we performed extensive functional studies for more than four years. As described in our method section, in fact, we used tag-free, tagged and endogenous *TMEM230*, and a large number of subcellular organelle markers for localization studies, including well known transmembrane proteins of synaptic vesicles, such as synaptophysin (SYN), vesicle associated membrane protein 2 (VAMP2) and vesicle monoamine transporter 2 (VMAT2). Regarding antibodies, all three anti-TMEM230 antibodies used in our report showed a single band with an expected size by Western blot. We hypothesized that TMEM230 is a transmembrane protein of synaptic vesicles primarily based on the following lines of strong evidence: (I) TMEM230 is transmembrane protein, it nicely co-localized with other well-known transmembrane proteins of synaptic vesicles (SYN, VAMP2 and VMAT2) (Fig. 1a-p; Supplementary Fig. 14-17); (II) TMEM230 was co-enriched in the synaptosome fraction, especially in the synaptic vesicle subfraction (Fig. 2q); (III) TMEM230 was clearly detected in the presynaptic vesicle pool region of the presynaptic axonal terminal by immunoGold electron microscopy (Fig. 2r). Apparently, TMEM230 may localizes to other secretory/recycling vesicles in other types of cells. However, there is no doubt that it predominantly localizes to synaptic vesicles in neurons, based on our data from primary neurons ^1^.

We understand that synaptic vesicles generally refer to the readily releasable pool in presynaptic axon terminals as we showed in Fig. 2r ^1^. However, synaptic vesicle precursors containing synaptic vesicle proteins, such as synaptophysin and VAMP2, are synthesized and preassembled in the cell body and transported to the axonal terminals, where they exert their physiological functions, and then they are recycled through endocytic mechanism. We used primary neuron culture system to monitor this vesicle trafficking. These vesicles were labeled by synaptic vesicle marker (eGFP-VAMP2). Because we were unable to classify these VAMP2-positive vesicles into specific synaptic vesicle pools, we did not specify the subpopulations they belong. Nonetheless, our purpose in this initial report was to identify possible defects related to synaptic vesicle trafficking in neurons when TMEM230 is mutated. The matured synaptic vesicle trafficking in the presynaptic pool could be better characterized using *in vivo* models and more comprehensive approaches in the future studies.

Dr. Farrer might have misunderstood our trafficking assay. In fact, wild-type and mutant TMEM230 proteins in our trafficking assay were not tagged. eGFP-VAMP2 was used to label synaptic vesicles. TMEM230 was expressed using pIRES2-ZsGreen1 vector. This vector simultaneously but independently expresses TMEM230 and ZsGreen. Because ZsGreen was distributed in a diffuse pattern in cells, it could be used not only to identify transfected cells, but also to select cells with a similar level of expression. However, we agree that using *in vivo* models would be a better approach. Our *in vivo* studies are currently in progress.

We respectfully disagree with Dr. Farrer at using the presumably non-pathogenic TMEM230 substitutions as controls. Because these variants are very rare (∼0.001) and their non-pathogenicity should not be presumed. We believe that the appropriate control is wild-type TMEM230, especially in the initial study.

**Comment-9.** We have studied the consequences of the *DNAJC13* c.2564A> G (p.Asn855Ser) mutation in synaptically mature primary cortical cultures (DIV21) prepared from a knock-in mouse (Figure 1b). We observe profound effects on endosomal tubulation in heterozygotes, and in endosomal proliferation in both heterozygous and homozygous mutant animals, with respect to neurons grown from wild-type embryos from the same litters. Our findings recapitulate and extend observations made in DNAJC13/RME-8 knockdown experiments in immortalized cell lines using the same SNX1-GFP endosome marker suggesting a gene-dose dependent ‘loss of function’ is induced by this mutation.

**Response-9.** We believe that the pathogenicity of *DNAJC13* mutations to PD should first derive from established genetic evidence. But such evidence is lacking for *DNAJC13* to date. Cell culture studies certainly advance the understanding of the mechanism of disease, only after the genetic cause has been firmly established. Mutations in many genes may disrupt cellular trafficking, that does not mean that they cause naturally-occurring PD. DNAJC13-N855S variant is not present in three (II-1, III-1 and III-23) out of 15 affected members of this family. We believe that this basically excludes this variant as disease-causing. However, TMEM230-R141L variant is present in 14 affected members, except for III-14 in this family. Moreover, we showed the other independent and unarguable genetic evidence, including two *TMEM230* stop codon variants in 10 familial PD patients in eight families with different ethnicities.

We appreciate the efforts by Dr. Farrer’s group to understand the function of DNAJC13. However, functional studies may not compensate for genetic evidence. Indeed, both TMEM230 and DNAJC13 appear to be involved in endosomal function, but multiple lines of strong genetic evidence support *TMEM230* as disease-causing. Such strong genetic evidence is lacking for *DNAJC13* in PD. We have recently shown evidence that TMEM230 plays an important role in Rab8a-mediated secretory vesicle trafficking and retromer trafficking, providing a potential mechanistic link to LRRK2, another PD-linked protein ^37^.

**Comment-10.** In conclusion, DNAJC13 c.2564A> G (p.Asn855Ser) remains compelling as the gene mutation for familial parkinsonism in this Mennonite kindred. Future research in DNAJC13 biology will undoubtedly make important contributions to the field, for which DNA constructs and the constitutive knock in mouse model are now available (on request). In contrast, the evidence TMEM230 is a gene for familial parkinsonism is not compelling. The mutations Deng and colleagues describe fail to segregate with disease, as claimed. TMEM230 p.Arg141Leu, at non-conserved amino acid residue, is also rather too frequent to be causal of disease. Neither is there any evidence to support the notion that DNAJC13 and TMEM230 mutations may act together, in concert, to lead to parkinsonism. We recommend Deng and colleagues reassess and retract their prior work. In the interim, we invite independent authors to take the opportunity to assess our genetic analysis and samples (Figure 1a), and publish their findings accordingly.

**Response-10.** As we responded above, in the original Mennonite PD family, DNAJC13-N855S mutation is not present in three affected individuals. In a total of 9,006 PD cases tested world-wide, only five single PD cases had this mutation, and no co-segregation of this mutation could be demonstrated with PD in any families. All these five single cases were found by Dr. Farrer’s group. Notably, all the DNAJC13-N855S-positive cases have Mennonite heritage (with the same haplotype). We believe that absence of the DNAJC13-N855S mutation in three affected individuals basically excludes the pathogenicity of this mutation to PD in this Mennonite PD family. Therefore, it is highly likely that the DNAJC13-N855S is a private variant specific to a Mennonite ethnic population, but not pathogenic for PD.

In contrast, TMEM230-R141L mutation was found in 14 affected individuals, with an exception to one affected member only (II-14). But considering 5% phenocopy rate of all affected PD individuals in the pedigrees shown by a previous study ^20^, this is quite a reasonable rate (1/15, 6.7%) in this family. The key supporting evidence is our identification of TMEM230-*184ProGlyext*5 mutation in nine PD patients in seven unrelated Chinese families. This mutation was shown to co-segregate with PD in two of the Chinese families. We also identified TMEM230-*184Trpext*5 mutation in a family PD patient in North America. Intriguingly, both mutations involve the stop codon, adding identical five amino acids (HPPHS) at the C terminus. We believe that identification of these two TMEM230 mutations (*184Trpext*5 and *184ProGlyext*5) alone provides solid genetic evidence for the pathogenicity of TMEM230 mutations to PD. Most recently, TMEM230-A110T mutation was identified in two unrelated PD cases, suggesting that TMEM230 may also be involved in the apparently sporadic PD cases ^38^.

Taken together, our new linkage data from genome-wide MegaEx chip screening containing ∼2 million SNPs demonstrated consistent high LOD scores in all models at the *TMEM230* locus. The *TMEM230* locus yielded the highest two-point and multi-point LOD scores in the affected-only model in the whole-genome. **No other loci in the entire genome yielded higher LOD scores than the chromosome 20p (TMEM230) locus in the affected-only model**. These new data are consistent with our previous linkage data using microsatellite markers. In addition, we re-analyzed our exome sequencing data and updated the clinical information (Table 1). TMEM230-R141L remains to be the best explanation for PD in this family, as this mutation can explain the affected status of the entire family with a single phenocopy (II-14). DNAJC13-N855S and ZBTB38-R264Q require three phenocopies, and the other mutations require five or more phenocopies (Table 1).

Overall, we remain confident that *TMEM230* mutations lead to PD, based on the genetic data from the original Mennonite PD family and the other eight families from North America and China, and recently obtained new data and information. This gene will provide an important clue and resource in dissecting the role of synaptic vesicle/endosomal trafficking/recycling in the pathogenesis of PD.

Based on our new clinical information, we updated and revised Supplementary Fig. 1, which is attached at the end of this response.

---

**Supplementary Fig. 1.**
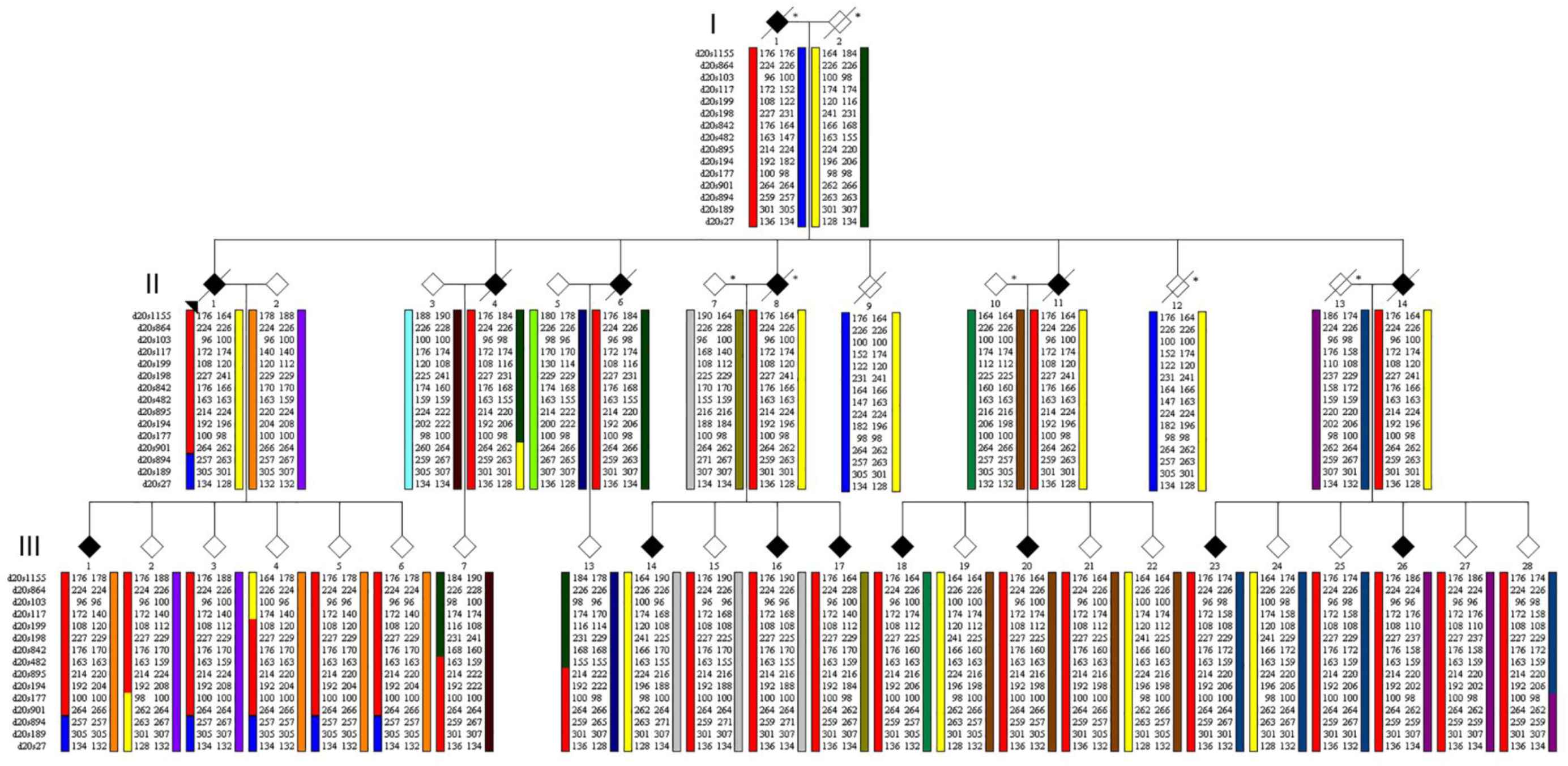
In the original article, supplementary figure 1 incorrectly assigned the disease status to III-15, instead of III-14. Supplementary figure 1 has now been revised. The authors apologize for this error. The authors remain confident that TMEM230-R141L is pathogenic, assuming a single phenocopy for individual III-14 in this large PD family.

